# Human electromagnetic and haemodynamic networks systematically converge in unimodal cortex and diverge in transmodal cortex

**DOI:** 10.1101/2021.09.07.458941

**Authors:** Golia Shafiei, Sylvain Baillet, Bratislav Misic

## Abstract

Whole-brain neural communication is typically estimated from statistical associations among electromagnetic or haemodynamic time-series. The relationship between functional network architectures recovered from these two types of neural activity remains unknown. Here we map electromagnetic networks (measured using magnetoencephalography; MEG) to haemodynamic networks (measured using functional magnetic resonance imaging; fMRI). We find that the relationship between the two modalities is regionally heterogeneous and systematically follows the cortical hierarchy, with close correspondence in unimodal cortex and poor correspondence in transmodal cortex. Comparison with the BigBrain histological atlas reveals that electromagnetic-haemodynamic coupling is driven by laminar differentiation and neuron density, suggesting that the mapping between the two modalities can be explained by cytoarchitectural variation. Importantly, haemodynamic connectivity cannot be explained by electromagnetic activity in a single frequency band, but rather arises from the mixing of multiple neurophysiological rhythms. Correspondence between the two is largely driven by MEG functional connectivity at the beta (15-29 Hz) frequency band. Collectively, these findings demonstrate highly organized but only partly overlapping patterns of connectivity in MEG and fMRI functional networks, opening fundamentally new avenues for studying the relationship between cortical microarchitecture and multi-modal connectivity patterns.

## INTRODUCTION

The structural wiring of the brain imparts a distinct signature on neuronal co-activation patterns. Inter-regional projections promote signaling and synchrony among distant neuronal populations, giving rise to coherent neural dynamics, measured as regional time series of electromagnetic or hemodynamic neural activity [46]. Systematic co-activation among pairs of regions can be used to map functional connectivity networks. Over the past decade, these dynamics are increasingly recorded without task instruction or stimulation; the resulting “intrinsic” functional connectivity is thought to reflect spontaneous neural activity.

The macro-scale functional architecture of the brain is commonly inferred from electromagnetic or haemodynamic activity. The former can be measured using electroencephalography (EEG) or magnetoencephalography (MEG), while the latter is measured using functional magnetic resonance imaging (fMRI). Numerous studies – using both MEG and fMRI – have reported evidence of intrinsic functional patterns that are highly organized [6, 12, 16, 19, 33, 107, 135, 148], reproducible [17, 29, 54, 101] and comparable to task-driven co-activation patterns [17, 30, 129].

How do electromagnetic and haemodynamic networks relate to one another? Although both modalities attempt to capture the same underlying biological process (neural activity), they are sensitive to different physiological mechanisms and ultimately reflect neural activity at fundamentally different time scales [5, 57, 62, 112, 113]. Emerging theories emphasize a hierarchy of time scales of intrinsic fluctuations across the cortex [50, 98, 110, 124], where unimodal cortex is more sensitive to immediate changes in the sensory environment, while transmodal cortex is more sensitive to prior context [7, 25, 26, 63, 72, 76]. This raises the possibility that the alignment between the relatively slower functional architecture captured by fMRI and faster functional architecture captured by MEG may systematically vary across the cortex.

Previous reports have found some, but not complete, global overlap between the two modalities. Multiple MEG and fMRI independent components – representing spatiotemporal signatures of resting-state intrinsic networks – show similar spatial topography, particularly the visual, somatomotor and default mode components [6, 16, 19, 70]. The spatial overlap between large-scale networks has also been reported in task-based studies and with networks recovered from other modalities, such as EEG and intracranial EEG [32, 45, 81, 91, 99]. Moreover, fMRI and MEG/EEG yield comparable fingerprinting accuracy, suggesting that they encode common information [31, 36, 44, 117]. Finally, global edge-wise comparisons between fMRI networks and electrocorticography (ECoG) [15], EEG [35, 146, 147] and MEG [51, 71, 134] also yield moderate correlations. Although global comparisons are more common when different modalities are studied, regional and network-level relationships have also been explored using electrophysiological and intracranial EGG recordings [32, 83, 97] as well as EEG and MEG recordings [71, 126, 131]. Regional comparisons of electrophysiological and fMRI recordings also suggest that the relationship between the two may be affected by distinct cytoarchitecture and laminar structure of brain regions, particularly in visual and frontal cortex [8, 10, 22, 84, 85, 120, 121, 127]. How the coupling between fMRI and MEG connectivity profiles varies from region to region, and how this coupling reflects cytoarchitecture, is still not fully understood. Furthermore, previous studies have mostly assessed the association between haemodynamic and electromagnetic networks for separate frequency bands, investigating independent contributions of individual rhythms to haemodynamic connectivity. This effectively precludes the possibility that superposition and mixing of elementary electromagnetic rhythms manifests as patterns of haemodynamic connectivity [71, 86, 134].

How regional connectivity profiles of MEG and fMRI functional networks are associated across the cortex and how their correspondence relates to the underlying cytoarchitecture, remains an exciting open question. Here, we use a linear multi-factor model that allows to represent the haemodynamic functional connectivity profile of a given brain region as a linear combination of its electromagnetic functional connectivity in multiple frequency bands. We then explore how the two modalities align across the neocortex and investigate the contribution of cytoarchitectonic variations to their alignment.

## RESULTS

Data were derived using task-free MEG and fMRI recordings in the same unrelated participants from the Human Connectome Project (HCP [140]; *n* = 33). We first develop a simple regression-based model to map regional MEG connectivity to regional fMRI connectivity using group-average data. We then investigate how regionally heterogeneous the correspondence between the two is, and how different rhythms contribute to this regional heterogeneity. Finally, we conduct extensive sensitivity testing to demonstrate that the results are robust to multiple methodological choices.

### Relating haemodynamic and electromagnetic connectivity

To relate fMRI and MEG functional connectivity patterns, we apply a multi-linear regression model [142] (Fig. 1). The model is specified for each brain region separately, attempting to predict a region’s haemodynamic connectivity profile from its electromagnetic connectivity profile. The dependent variable is a row of the fMRI functional connectivity (FC) matrix and the independent variables are the corresponding rows of MEG FC matrices for six canonical electrophysiological bands, estimated using amplitude envelope correlation (AEC [20]) with spatial leakage correction (See “Methods” for more details). For a model fitted for a given node *i*, the observations in the model are the connections of node *i* to the other *j* ≠ *i* regions (Fig. 1a). The model predicts the fMRI FC profile of node *i* (i.e. *i*-th row) from a linear combination of MEG FC profiles of node *i* in the six frequency bands (i.e. *i*-th rows of MEG FC matrices). Collectively, the model embodies the idea that multiple rhythms could be superimposed to give rise to regionally heterogeneous haemodynamic connectivity.

**Figure 1.**
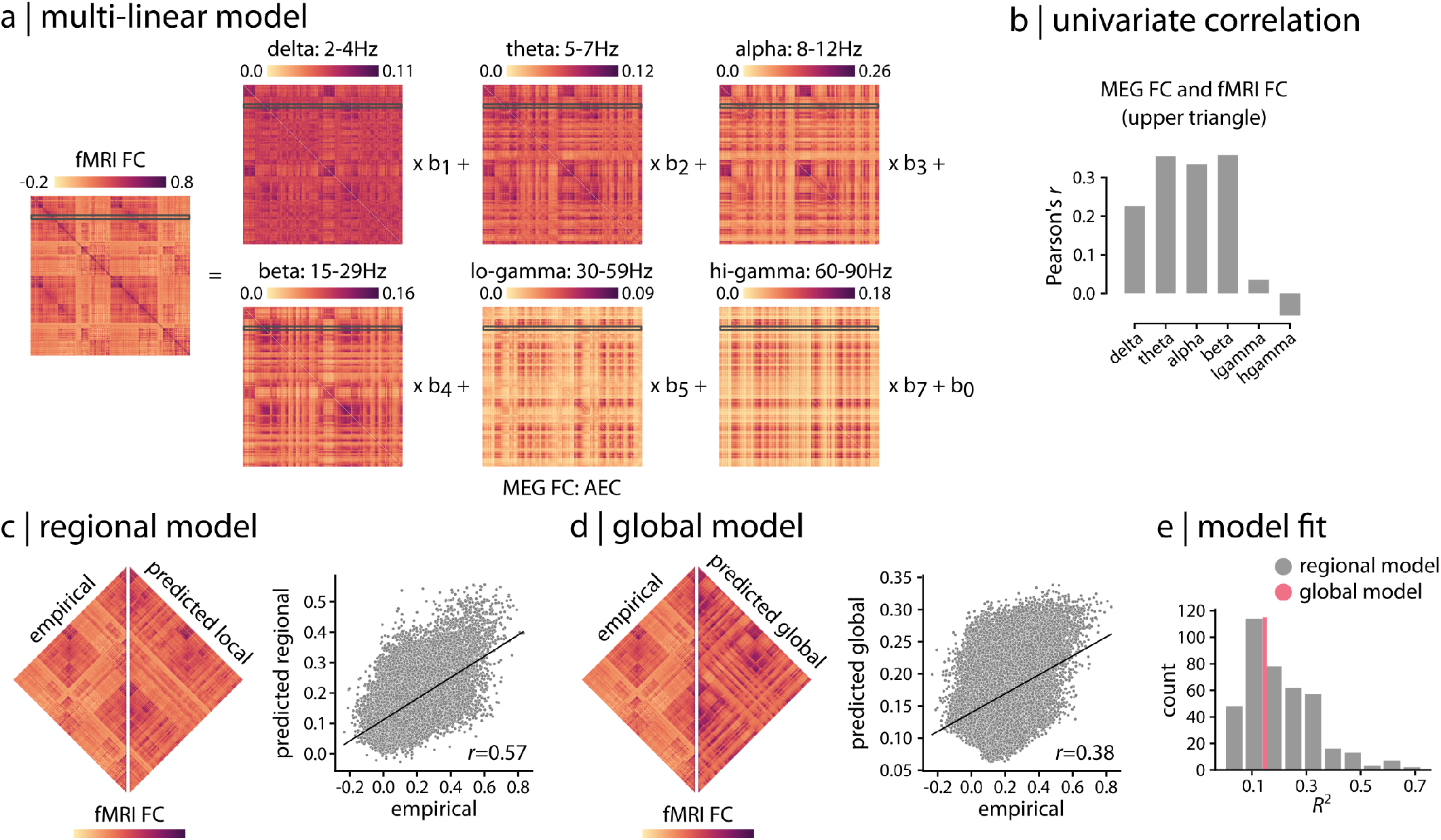
Relating haemodynamic and electromagnetic connectivity. (a) A multi-linear regression model was applied to predict resting state fMRI connectivity patterns from band-limited MEG functional connectivity (amplitude envelope correlation; AEC [20]). The model is specified for each brain region separately, attempting to predict a region’s haemodynamic connectivity profile from its electromagnetic connectivity profile. (b) The overall relationship between fMRI and MEG functional connectivity is estimated by correlating the upper triangle of fMRI FC (i.e. above diagonal) with the upper triangles of band-limited MEG FC, suggesting moderate relationship between the two across frequency bands. (c) Regional multi-linear model shown in panel (a) is used to predict fMRI FC from band-limited MEG FC for each brain region (i.e. row) separately. The empirical and predicted fMRI FC are depicted side-by-side for the regional model. The whole-brain edge-wise relationship between the empirical and predicted values is shown in the scatter plot. Each grey dot represents an edge (pairwise functional connection) from the upper triangles of empirical and predicted fMRI FC matrices. (d) A global multi-linear model is used to predict the entire upper triangle of fMRI FC from the upper triangles of the MEG FC matrices. The empirical and predicted fMRI FC are depicted side-by-side for the global model. The whole-brain edge-wise relationship between the empirical and predicted values is shown in the scatter plot. Each grey dot represents en edge from the upper triangles of empirical and predicted fMRI FC matrices. (e) The distribution of regional model fit quantified by *R*^2^ is shown for regional model (grey histogram plot). The global model fit is also depicted for comparison (pink line). The data and code needed to generate this figure can be found in https://github.com/netneurolab/shafiei_megfmrimapping and https://zenodo.org/record/6728338.

Indeed, we find that the relationship between haemodynamic and electromagnetic connectivity is highly heterogeneous. Band-limited MEG connectivity matrices are moderately correlated with fMRI connectivity, ranging from *r* = −0.06 to *r* = 0.36 (Fig. 1b; *r* denotes Pearson correlation coefficient). The regional multi-linear model fits range from adjusted-*R*^2^ = −0.002 to adjusted-*R*^2^ = 0.72 (*R*^2^ denotes coefficient of determination; hereafter we refer to adjusted-*R*^2^ as *R*^2^), suggesting a close correspondence in some regions and poor correspondence in others (Fig. 1c,e). Band-specific regional model fits are depicted in Fig. S1, where each band-specific MEG connectivity is separately used as a single predictor in the model. For comparison, a single global model is fitted to the data, predicting the entire upper triangle of the fMRI FC matrix (i.e. all values above the diagonal) from a linear combination of the upper triangles of six MEG FC matrices (i.e. all values above the diagonal)(See “Methods” for more detail). The global model, which simultaneously relates whole-brain fMRI FC to the whole-brain MEG FC, yields an *R*^2^ = 0.15 (Fig. 1d,e). Importantly, the global model clearly obscures the wide range of correspondences, which can be considerably greater or smaller for individual regions.

### Hierarchical organization of cross-modal correspondence

We next consider the spatial organization of fMRI-MEG correspondence. Fig. 2a shows the spatial distribution of regional *R*^2^ values, representing regions with low or high correspondence. Regions with strong cross-modal correspondence include the visual, somato-motor and auditory cortex. Regions with low cross-modal correspondence include the posterior cingulate, lateral temporal and medial prefrontal cortex.

**Figure 2.**
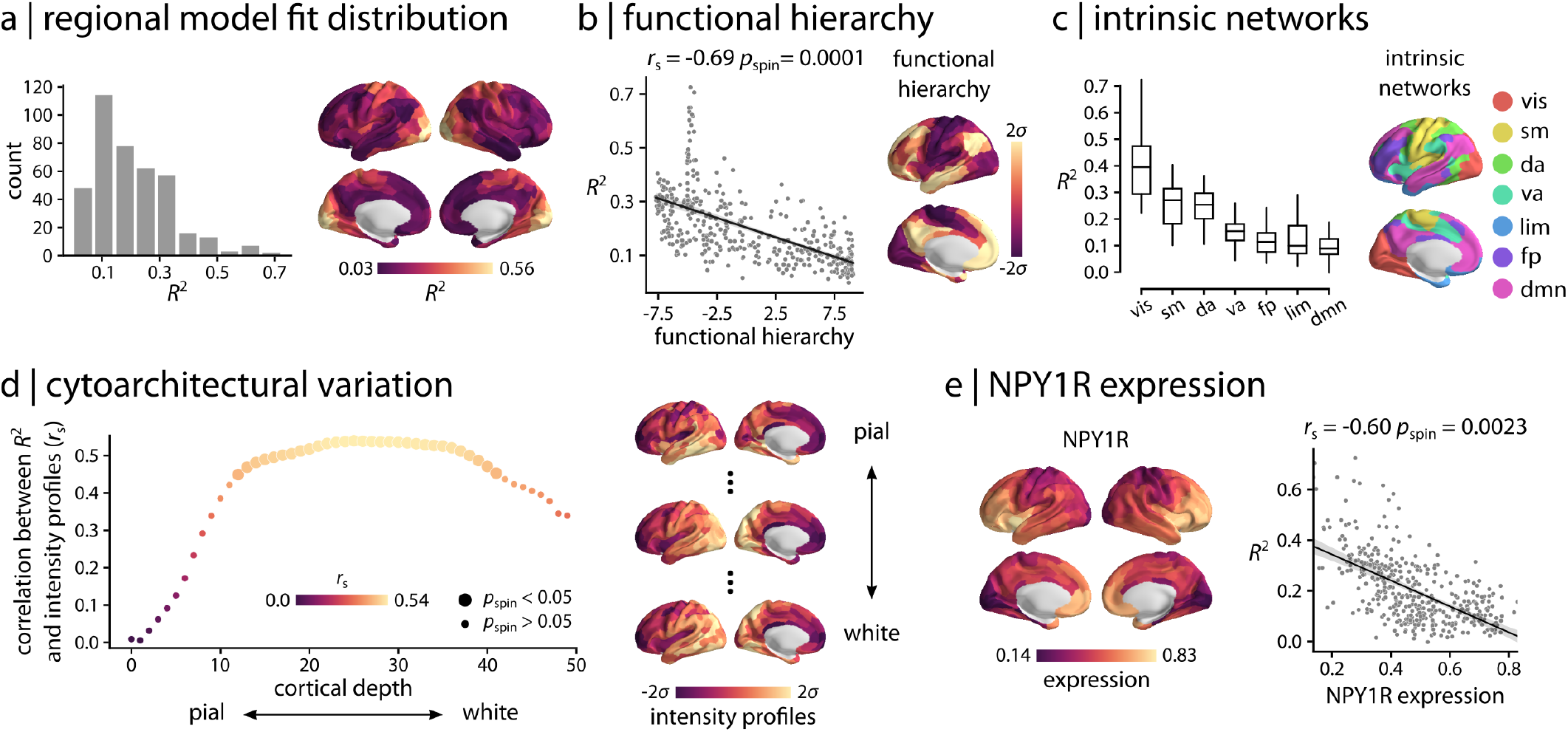
Regional model fit. (a) Spatial organization of fMRI-MEG correspondence is depicted for the regional model fit (95% interval). The cross-modal correspondence of connectivity profiles of brain regions is distributed heterogeneously across the cortex, representing regions with low or high correspondence. Strong cross-modal correspondence is observed in sensory areas whereas poor correspondence is observed for higher order regions. (b) Spatial organization of the cross-modal correspondence is compared with the functional hierarchical organization of cerebral cortex [87]. The two are significantly anti-correlated, confirming poor fMRI-MEG correspondence in connectivity profile of higher-order, transmodal areas compared to strong correspondence for sensory, unimodal regions. (c) Regions are stratified by their affiliation with macro-scale intrinsic networks [148]. The distribution of *R*^2^ is depicted for each network, displaying a systematic gradient of cross-modal correspondence with the highest correspondence in the visual network and lowest correspondence in the default mode network. (d) The model fit is related to the cytoarchitectural variation of the cortex, estimated from the cell staining intensity profiles at various cortical depths obtained from the BigBrain histological atlas [2, 104]. Bigger circles denote statistically significant associations after correction for multiple comparisons by controlling the false discovery rate (FDR) at 5% alpha [13]. The peak association between cross-modal correspondence and cytoarchitecture is observed approximately at cortical layer IV that has high density of granule cells. Staining intensity profiles are depicted across the cortex for the most pial, the middle and the white matter surfaces. (e) Microarray gene expression of vasoconstrictive NPY1R (Neuropeptide Y Receptor Y1) was estimated from the Allen Human Brain Atlas (AHBA; [69]). The MEG-fMRI cross-modal correspondence *R*^2^ map (i.e. regional model fit) is compared with NPY1R gene expression. *r*_s_ denotes Spearman rank correlation. Intrinsic networks: vis = visual; sm = somatomotor; da = dorsal attention; va = ventral attention; lim = limbic; fp = frontoparietal; dmn = default mode. The data and code needed to generate this figure can be found in https://github.com/netneurolab/shafiei_megfmrimapping and https://zenodo.org/record/6728338.

Collectively, the spatial layout of cross-modal correspondence bears a resemblance to the unimodal-transmodal cortical hierarchy observed in large-scale functional and microstructural organization of the cerebral cortex [76]. To assess this hypothesis, we first compared the cross-modal *R*^2^ map with the principal functional hierarchical organization of the cortex, estimated using diffusion map embedding [80, 87] (Fig. 2b; see “Methods” for more details). The two are significantly anti-correlated (Spearman rank correlation coefficient *r*_s_ = −0.69, *p*_spin_ = 0.0001), suggesting strong cross-modal correspondence in unimodal sensory cortex and poor correspondence in transmodal cortex. We then stratify regions by their affiliation with macro-scale intrinsic networks and computed the mean *R*^2^ in each network [148] (Fig. 2c). Here we also observe a systematic gradient of cross-modal correspondence, with the strongest correspondence in the visual network and poorest correspondence in the default mode network.

We relate the cross-modal *R*^2^ map to the cytoarchitectural variation of the cortex (Fig. 2d). We use the Big-Brain histological atlas to estimate granular cell density at multiple cortical depths [2, 104]. Cell-staining intensity profiles were sampled across 50 equivolumetric surfaces from the pial surface to the white matter surface to estimate laminar variation in neuronal density and soma size. Fig. 2d shows the correlation between MEG-fMRI correspondence and cell density (*y*-axis) at different cortical depths (*x*-axis). Interestingly, the model fit is associated with cytoarchitectural variation of the cortex, with the peak association observed approximately at cortical layer IV that has high density of granular cells and separates supra-and infra-granular layers [102, 103, 144]. Layer IV predominately receives feedforward projections and has high vascular density [39, 61, 122]. We further assess the relationship between MEG-fMRI crossmodal correspondence and vascular attributes. We obtain the microarray gene expression of the vasoconstrictive NPY1R (Neuropeptide Y Receptor Y1) from Allen Human Brain Atlas (AHBA; [69]; see “Methods” for more details), given previous reports that the BOLD response is associated with the vasoconstrictive mechanism of Neuropeptide Y (NPY) acting on Y1 receptors [138]. We then compare the cross-modal association map with the expression of NPY1R and identify a significant association between the two (Fig. 2e; *r*_s_ = −0.60, *p*_spin_ = 0.0023). This demonstrates that regions with low cross-modal correspondence are enriched for NPY1R whereas areas with high cross-modal associations have less NPY-dependent vasoconstriction. Altogether, the results suggest that the greater coupling in unimodal cortex may be driven by the underlying cytoarchitecture, reflecting higher density of granular cells and distinct vascularization of cortical layer IV.

We also relate cross-modal *R*^2^ map to the variation of structure-function coupling across the cortex, which has also been shown to follow the unimodal-transmodal hierarchy [11, 108, 132, 142, 150]. We estimate structurefunction coupling as the Spearman rank correlation between regional structural and functional connectivity profiles [11] (Fig. S2; see “Methods” for more details). We then correlate the identified map with the regional model fit, identifying a significant association between the two (Fig. S2; *r*_s_ = 0.40, *p*_spin_ = 0.0025). This is consistent with the notion that both haemodynamic and electromagnetic neural activity are constrained by the anatomical pathways and the underlying structural organization [24, 118, 130].

### Heterogeneous contributions of multiple rhythms

How do different rhythms contribute to regional patterns of cross-modal correspondence? To address this question and to assess the effects of cross-correlation between MEG connectivity at different frequency bands (Fig. S5), we perform a dominance analysis for every regional multi-linear model [4, 21]. Specifically, dominance analysis is used to examine the separate effects of each band-limited MEG functional connectivity, as well as the effects of all other possible combinations of bandlimited MEG FC, on the regional model fit. This technique estimates the relative importance of predictors by constructing all possible combinations of predictors and re-fitting the multi-linear model for each combination. The possible combinations of predictors include sets of single predictors, all possible pairs of predictors, all possible combinations with 3 predictors, and so on. To assess the influence of each band on the model fit, dominance analysis re-fits the model for each combination and quantifies the relative contribution of each predictor as the increase in variance explained after adding that predictor to the models (i.e. gain in adjusted-*R*^2^). Fig. 3a shows the global dominance of each frequency band, where dominance is quantified as “percent relative importance” or “contribution percentage” of each band. Overall, we observe the greatest contributions from MEG connectivity at beta band, followed by theta and alpha bands, and smallest contributions from low and high gamma bands.

**Figure 3.**
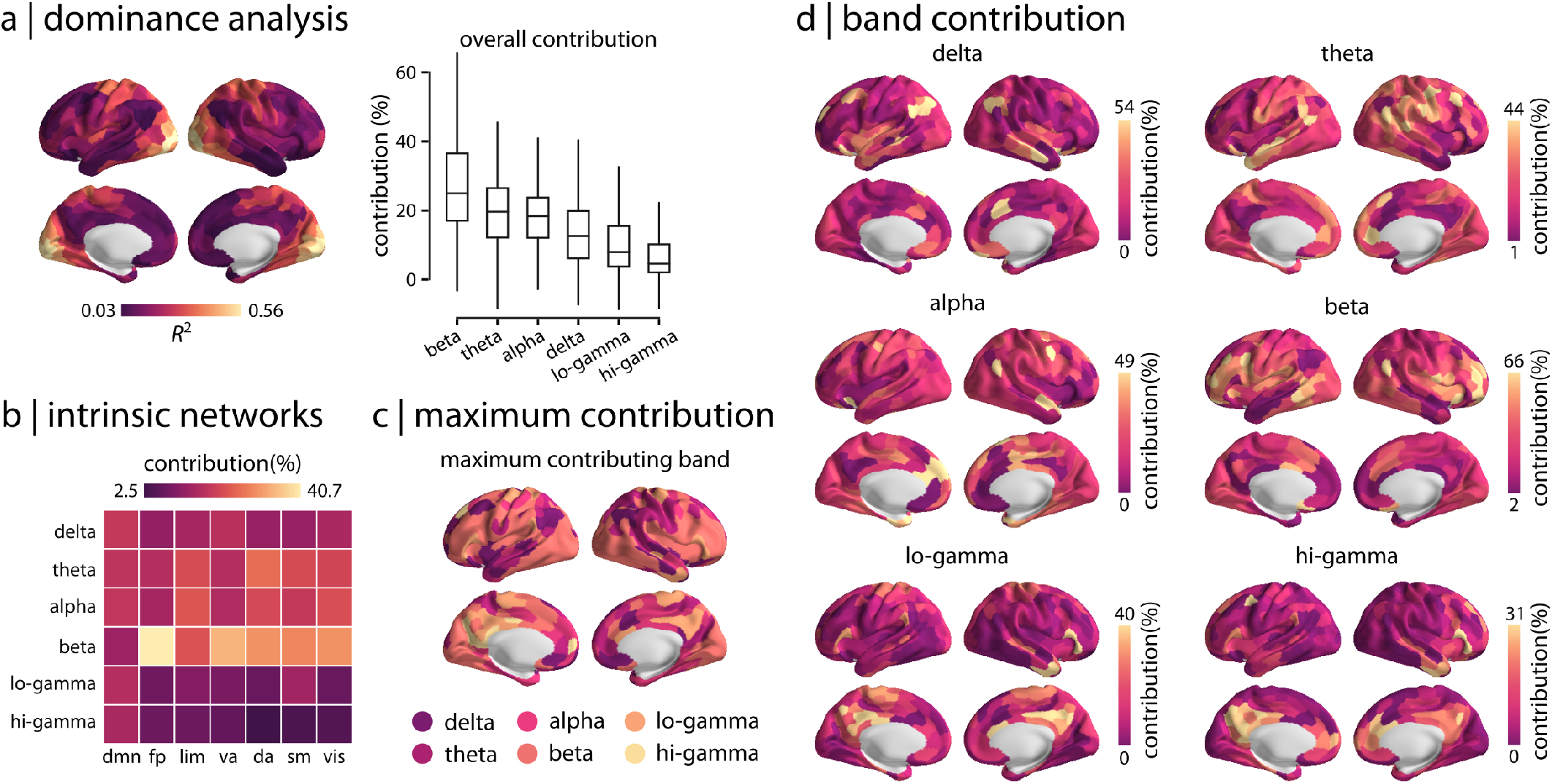
Dominance analysis. Dominance analysis is performed for each regional multi-linear model to quantify how MEG connectivity at different rhythms contribute to regional patterns of cross-modal correspondence [4, 21]. (a) The overall contribution of each frequency band is depicted for the regional model (box plots). Beta band connectivity, followed by theta and alpha bands, contribute the most to the model fit whereas low and high gamma bands contribute the least. (b) The mean contribution of different rhythms is estimated for the intrinsic networks. Consistent with the overall contributions depicted in panel (a), the greatest contribution is associated with beta band connectivity. (c) The most dominant predictor (frequency band) is depicted for each brain region, confirming overall higher contributions from beta band across the cortex. (d) Frequency band contribution to the regional cross-modal correspondence is shown separately for different rhythms across the cortex (95% intervals). The data and code needed to generate this figure can be found in https://github.com/netneurolab/shafiei_megfmrimapping and https://zenodo.org/record/6728338.

Zooming in on individual regions and intrinsic networks, we find that the dominance pattern is also regionally heterogeneous. Namely, the make-up and contribution of specific MEG frequencies to a region’s fMRI connectivity profile varies from region to region. Fig. 3b shows the dominance of specific rhythms in each intrinsic network. Fig. 3c shows the most dominant predictor for every brain region. We find that beta band contribution is highest in occipital and lateral frontal cortices. Sensorimotor cortex has high contributions from combinations of beta, alpha, and theta bands. Parietal and temporal areas are mostly dominated by delta and theta bands as well as some contribution from alpha band. Medial frontal cortex shows contributions from the alpha band, while low and high gamma bands contribute to posterior cingulate cortex and precuneus. Fig. 3d shows the dominance of specific rhythms separately for each region. Overall, we observe that beta connectivity has the highest contribution percentage (95% confidence interval: [2% 66%]), largely contributing to model prediction across the cortex. These findings are consistent with previous reports, demonstrating that haemodynamic connectivity is related to the superposition of band-limited electromagnetic connectivity and that band contributions vary across the cortex [71, 134].

Finally, we used Analysis of Variance (ANOVA) to quantitatively assess the differences in band-specific contributions to the cross-modal correspondence map (Table. S1). Specifically, we assessed the significance and effect size of differences in band-specific contributions for all possible pairs of frequency bands. We identify two main findings (for full results see Table. S1): (1) Overall, the variability of band-specific contributions is significantly larger between groups (i.e. bands) compared to the variability within groups (*F* (5, 2394) = 117.31; *p* < 0.0001). (2) Band-specific contributions are significantly different from each other and are ranked in the same order as depicted in Fig. 3a. Specifically, contribution of beta band is significantly larger than contribution of alpha band (difference of the means = 8.65, *t*-value = 9.46, *p*-value < 0.0001, Cohen’s *d* = 0.69) and theta band (difference of the means = 7.56, *t*-value = 8.27, *p*-value < 0.0001, Cohen’s *d* = 0.58). Also, the contribution from the delta band is significantly lower than beta (difference of the means = 12.37, *t*-value = 13.53, *p*-value < 0.0001, Cohen’s *d* = 0.96), alpha (difference of the means = 3.72, *t*-value = 4.07, *p*-value = 0.0007, Cohen’s *d* = 0.29), and theta (difference of the means = 4.81, *t*-value = 5.26, *p*-value < 0.0001, Cohen’s *d* = 0.37). Note that although the difference between alpha and theta band contributions is not significant, both their contributions are significantly lower than beta band and larger than delta band. Moreover, delta band contribution is significantly larger than contribution of lo-gamma (difference of the means = 3.78, *t*-value = 4.14, *p*-value = 0.0005, Cohen’s *d* = 0.29) and lo-gamma contribution is significantly larger than hi-gamma (difference of the means = 3.72, *t*-value = 4.07, *p*-value = 0.0007, Cohen’s *d* = 0.29). Note that the values reported here are the absolute values for difference of the means, *t*-values, *p*-values and Cohen’s *d* (effect size). All *p*-values are corrected for multiple comparisons using Bonferroni correction.

### Sensitivity analysis

Finally, we note that the present report goes through several decision points that have equally-justified alternatives. Here we explore the other possible choices. First, rather than framing the report from an explanatory perspective (focusing on model fit), we instead derive an equivalent set of results using a predictive perspective (focusing on out-of-sample prediction). We perform cross-validation at both the region-and subject-level (Fig. 4a,b). For region-level cross-validation, we pseudorandomly split the connectivity profile of a given region into train and test sets based on spatial separation (inter-regional Euclidean distance), such that 75% of the closest regions to a random region are selected as the train set and the remaining 25% of the regions are selected as test set (399 repetitions; see “Methods” for more details) [59]. We then train the multi-linear model using the train set and predict the connection strength of the test set for each region and each split. The mean regional model performance across splits is consistent for train and test sets (Fig. 4a; *r* = 0.78, *p*_spin_ = 0.0001). For subject-level cross-validation, we use leave-one-out-cross validation, wherein we train the regional multi-linear models using data from *n* −1 subjects and test each one on the held-out subject. The mean regional model performance is consistent for train and test sets (Fig. 4b; *r* = 0.90, *p*_spin_ = 0.0001). Altogether, both analyses give similar, highly concordant results with the simpler model fit-based analysis, identifying strong cross-modal correspondence in unimodal sensory regions and poor correspondence in transmodal areas.

**Figure 4.**
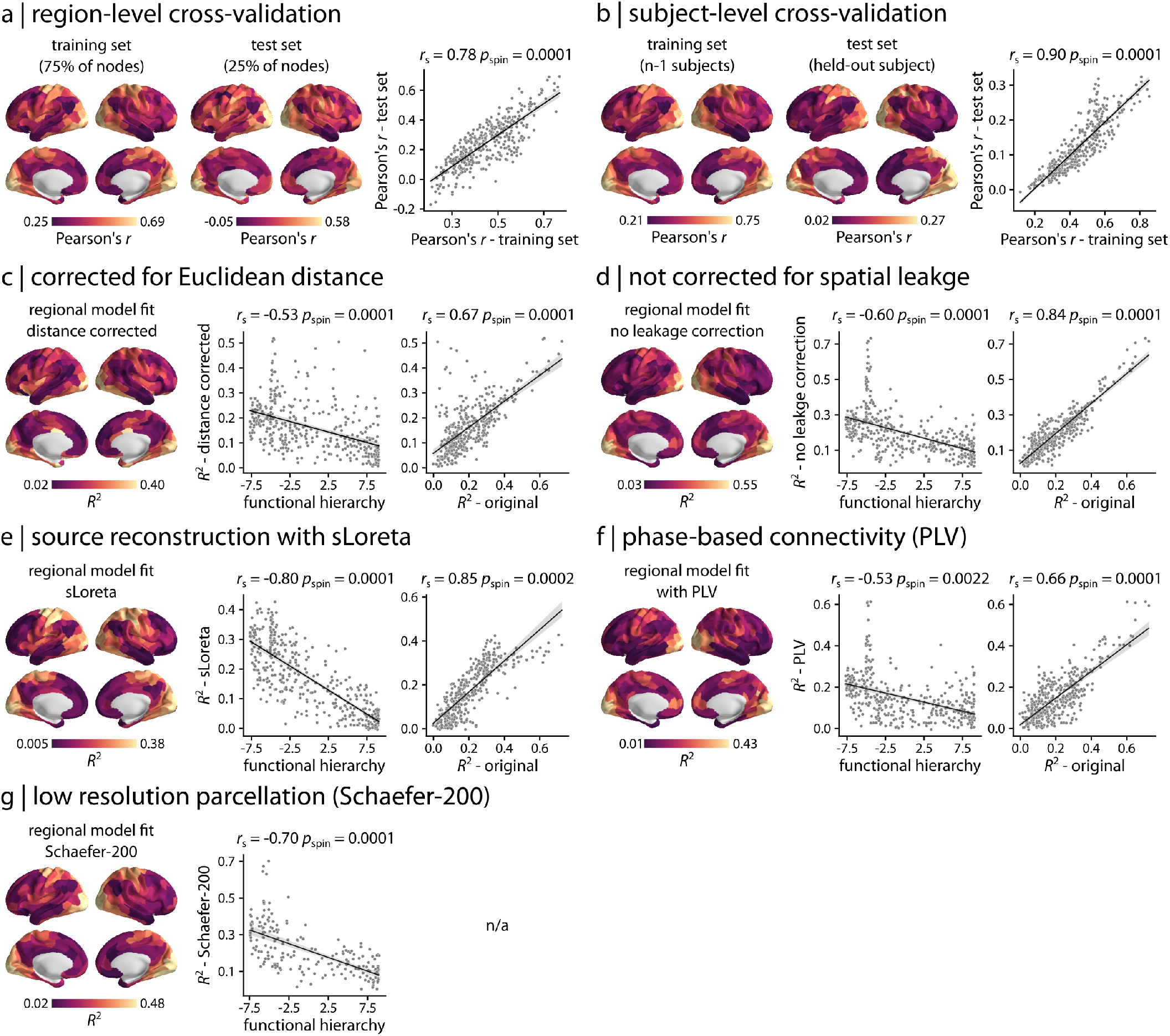
Sensitivity analysis. (a) A regional cross-validation was performed by pseudorandomly splitting the connectivity profile of a given region into train and test sets based on spatial separation (See “Methods” for more details). The multi-linear model is then fitted on the train set and is used to predict the connection strength of the test set for each region and each split. The mean regional model performance across splits is depicted for train and test sets, displaying consistent results between the two (scatter plot). The out-of-sample model performance is stronger in the sensory, unimodal areas compared to transmodal areas, consistent with original findings (Fig. 2). (b) A subject-level cross-validation was performed using a leave-one-out approach. The regional multi-linear model is trained using data from *n* −1 subjects and is tested on the held-out subject for each region separately. The mean regional model performance is shown for train and test sets, displaying consistent results between the two (scatter plot). The out-of-sample model performance is stronger in the sensory, unimodal areas compared to transmodal areas, consistent with original findings (Fig. 2). The analysis is also repeated for various processing choices: (c) after regressing out interregional Euclidean distance from connectivity matrices, (d) using MEG connectivity data without spatial leakage correction, (e) using another MEG source reconstruction method (standardized low resolution brain electromagnetic tomography; sLoreta [105]), (f) using a phase-based MEG connectivity measure (phase-locking value; PLV [79, 96]), and (g) at a low resolution parcellation (Schaefer-200 atlas [119]). The results are consistent across all control analyses, identifying similar cross-modal correspondence maps as the original analysis (Fig. 2a). All brain maps are shown at 95% intervals. *r*_s_ denotes Spearman rank correlation. The data and code needed to generate this figure can be found in https://github.com/netneurolab/shafiei_megfmrimapping and https://zenodo.org/record/6728338.

To consider the effect of spatial proximity on the findings, we remove the exponential inter-regional Euclidean distance trend from all connectivity matrices before fitting any model. The results are consistent with and without distance correction (Fig. 4c; correlation with functional hierarchy: *r*_s_ = −0.53, *p*_spin_ = 0.0001; correlation with original *R*^2^: *r*_s_ = 0.67, *p*_spin_ = 0.0001). We also obtain consistent findings when we repeat the analysis without accounting for spatial leakage effect in estimating MEG connectivity with AEC (Fig. 4d; correlation with functional hierarchy: *r*_s_ = −0.60, *p*_spin_ = 0.0001; correlation with original *R*^2^: *r*_s_ = 0.84, *p*_spin_ = 0.0001). Next, we use another source reconstruction method (standardized low resolution brain electromagnetic tomography; sLoreta [105]) instead of LCMV beamformers, as previous reports suggest that sLoreta improves source localization accuracy [65, 67]. We then estimate MEG connectivity with AEC and repeat the multi-linear model analysis, identifying similar results as before (Fig. 4e; correlation with functional hierarchy: *r*_s_ = −0.80, *p*_spin_ = 0.0001; correlation with original *R*^2^: *r*_s_ = 0.85, *p*_spin_ = 0.0002). Next, we compute MEG connectivity using an alternative, phase-based connectivity measure (phase locking value; PLV [79, 96]), rather than the AEC. The two FC measures yield similar cross-modal correspondence maps (Fig. 4f; correlation with functional hierarchy: *r*_s_ = −0.53, *p*_spin_ = 0.0022; correlation with original *R*^2^: *r*_s_ = 0.66, *p*_spin_ = 0.0001). We also repeat the analysis using a low resolution parcellation (Schaefer-200 atlas [119]) to ensure that the findings are independent from the choice of parcellation. As before, the results demonstrate similar cross-modal correspondence map (Fig. 4g; correlation with functional hierarchy: *r*_s_ = −0.70, *p*_spin_ = 0.0001). To assess the extent to which the results are influenced by MEG source localization error, we compare the cross-modal correspondence pattern to peak localization error estimated using crosstalk function (CTF) [64, 65, 67, 82, 95]. No significant association is observed between *R*^2^ pattern and CTF for LCMV (Fig. S3a; *r*_s_ = −0.14, *p*_spin_ = 0.6) and sLoreta (Fig. S3b; *r*_s_ = −0.04, *p*_spin_ = 0.9) source reconstruction solutions. Finally, to confirm that the cross-modal correspondence pattern is independent from signal-to-noise ratio (SNR), we compare the regional model fit with the SNR map of the reconstructed sources, identifying no significant association between the two (Fig. S4; *r*_s_ = 0.32, *p*_spin_ = 0.25)(See “Methods” for more details).

## DISCUSSION

In the present report we map electromagnetic functional networks to haemodynamic functional networks in the human brain. We find two principal results. First, the relationship between the two modalities is regionally heterogeneous but systematic, reflecting the unimodal-transmodal cortical hierarchy and cytoarchitectural variation. Second, haemodynamic connectivity cannot be explained by electromagnetic connectivity in a single band, but rather reflects mixing and superposition of multiple rhythms.

The fact that the association between the two modalities follows a gradient from unimodal to transmodal cortex resonates with emerging work on cortical hierarchies [76, 87, 93]. Indeed, similar spatial variations are observed for multiple micro-architectural features, such as gene expression [23, 49, 59], T1w/T2w ratio [75], laminar differentiation [144] and neurotransmitter receptor profiles [47, 55, 60]. Collectively, these studies point to a natural axis of cortical organization that encompasses variations in both structure and function across micro-, meso- and macro-scopic spatial scales.

Interestingly, we find the closest correspondence between fMRI and MEG functional connectivity in unimodal cortex (including the visual and somatomotor networks) and the poorest correspondence in transmodal cortex (default mode, limbic, fronto-parietal and ventral attention networks). In other words, the functional architectures of the two modalities are consistent early in the cortical hierarchy, presumably reflecting activity related to instantaneous changes in the external environment. Conversely, as we move up the hierarchy, there is a gradual separation between the two architectures, suggesting that they are differently modulated by endogenous inputs and contextual information. How the two types of functional connectivity are related to ongoing task demand is an exciting question for future research.

Why is there systematic divergence between the two modalities? Our findings suggest that topographic variation in MEG-fMRI coupling is due to variation in cytoarchitecture and neurovascular coupling. First, we observe greater MEG-fMRI coupling in regions with prominent granular layer IV. This result may reflect variation of microvascular density at different cortical layers [40, 120, 122]. Namely, cortical layer IV is the most vascularized, and this is particularly prominent in primary sensory areas [122]. The BOLD response mainly reflects local field potentials arising from synaptic currents of feedforward input signals to cortical layer IV [39, 61]; as a result, the BOLD response is more sensitive to cortical layer IV with high vascular density [139]. Therefore, electromagnetic neuronal activity originating from layer IV should be accompanied by a faster and more prominent BOLD response. This is consistent with our finding that brain regions with more prominent granular layer IV (i.e. unimodal cortex) have greater correspondence between electromagnetic and haemodynamic functional architectures. In other words, heterogeneous cortical patterning of MEG-fMRI coupling may reflect heterogeneous patterning of underlying neurovascular coupling.

Second, we observe prominent anticorrelations between vasoconstrictive NPY1R-expressing neurons and MEG-fMRI coupling. Multiple studies of vasodilator and vasoconstrictor mechanisms involved in neural signaling have demonstrated links between microvasculature and the BOLD signal [40, 138]. For example, an optogenetic and 2-photon mouse imaging study found that task-related negative BOLD signal is mainly associated with vasoconstrictive mechanism of Neuropeptide Y (NPY) acting on Y1 receptors, suggesting that neurovascular coupling is cell specific [138]. Interestingly, by comparing the cortical expression of NPY1R (Neuropeptide Y Receptor Y1) in the human brain with MEG-fMRI correspondence pattern identified here, we find that regions with low cross-modal correspondence are enriched for NPY1R whereas areas with high cross-modal associations have less NPY-dependent vasoconstriction. Collectively, these results suggest that MEG-fMRI correspondence is at least partly due to regional variation in cytoarchitecture and neurovascular coupling.

More generally, numerous studies have investigated the laminar origin of cortical rhythms. For example, animal electrophysiological recordings demonstrated that visual and frontal cortex gamma activity can be localized to superficial cortical layers (supragranular layers I-III and granular layer IV), whereas alpha and beta activity are localized to deep infragranular layers (layers V-IV) [8, 10, 22, 84, 85, 127]. Similar findings have been reported in humans using EEG and laminar-resolved BOLD recordings, demonstrating that gamma and beta band EEG power are associated with superficial and deep layer BOLD response, respectively, whereas alpha band EEG power is associated with BOLD response in both superficial and deep layers [121]. Laminar specificity of cortical rhythms is increasingly emphasized in contemporary accounts of predictive processing [9]. In the predictive coding framework, transmodal regions generate predictive signals that modulate the activity of sensory unimodal regions depending on context [38]. These top-down signals are relatively slow, as they evolve with the context of exogenous (stimulation) inputs. The consequence on unimodal areas is a boost of their encoding gain, reflected in stronger, faster activity that tracks incoming stimuli. They in turn generate error signals that are slower and reflect the discrepancy between the predictions received and the actual external input. These slower error signals are then registered by higher-order transmodal regions. Specific cortical layers and rhythms contribute to this predictive coding [9]. For example, an unfamiliar, unpredicted stimulus is associated with increased gamma power that is fed forward up the cortical hierarchy (i.e. bottom-up from sensory to association cortices) through the superficial layers to transfer the prediction errors. This in turn results in low topdown, feedback predictions through deep cortical layers via alpha and beta rhythms. Conversely, predicted stimuli are associated with stronger feedback alpha and beta rhythms via deep layers, inhibiting the gamma activity for expected exogenous inputs [9]. This hierarchical predictive processing framework is also thought to underlie conscious perception by top-down transfer of perceptual predictions via alpha and beta rhythms through deep layers and bottom-up transfer of prediction errors via gamma rhythm through superficial layers, minimizing predictions errors [9, 114, 123]. Our results, linking cytoarchitecture with rhythm-specific connectivity may help to further refine and develop this emerging framework.

Altogether, our findings suggest that the systemic divergence between MEG and fMRI connectivity patterns may reflect variations in cortical cytoarchitecture and vascular density of cortical layers. However, note that due to the low spatial resolution of fMRI and MEG data, haemodynamic and electromagnetic connectivity is not resolved at the level of cortical layers. Rather, comparisons with cytoarchitecture are made via proxy datasets, such as the BigBrain histological atlas [2] and the Allen Human Brain Atlas [69]. Future work is required to assess the laminar-specificity of the cross-modal association in a more direct and comprehensive manner [41, 42, 73, 74].

Throughout the present report, we find that fMRI networks are best explained as arising from the superposition of multiple band-limited MEG networks. Although previous work has focused on directly correlating fMRI with MEG/EEG networks in specific bands, we show that synchronized oscillations in multiple bands could potentially combine to give rise to the well studied fMRI functional networks. Indeed, and as expected, the correlation between any individual band-specific MEG network and fMRI is substantially smaller than the multi-linear model that takes into account all bands simultaneously. Previous work on cross-frequency interactions [43] and on multi-layer MEG network organization [18] has sought to characterize the participation of individual brain regions within and between multiple frequency networks. Our findings build on this literature, showing that the superimposed representation may additionally help to unlock the link between MEG and fMRI networks.

It is noteworthy that the greatest contributions to the link between the two modalities came from beta band connectivity. Multiple authors have reported that – since it captures slow haemodynamic co-activation – fMRI network connectivity would be mainly driven by slower rhythms [19, 35, 43, 81, 86, 112]. Our findings demonstrate that although all frequency bands contribute to the emergence of fMRI networks, the greatest contributions come from beta band connectivity, followed by theta and alpha connectivity.

The present results raise two important questions for future work. First, how does structural connectivity shape fMRI and MEG functional networks [24, 132, 147]? We find that cross-modal correspondence between MEG and fMRI functional networks is associated with structure-function coupling measured from MRI functional and structural connectivity networks, suggesting that the cross-modal map may be constrained by structural connectivity. Previous reports demonstrate that unimodal, sensory regions have lower neural flexibility compared to transmodal, association areas and are more stable during development and evolution [115, 124, 149]. This suggests that the underlying anatomical network constrains neural activity and functional flexibility in a nonuniform manner across the cortex, resulting in higher degrees of freedom in structure-function coupling in regions related to highly flexible cognitive processes. However, given that MEG and fMRI capture distinct neurophysiological mechanisms, it is possible that haemodynamic and electromagnetic architectures have a different relationship with structural connectivity and this could potentially explain why they systematically diverge through the cortical hierarchy [11, 108, 132, 142, 150]. Second, the present results show how the two modalities are related in a task-free resting state, but what is the relationship between fMRI and MEG connectivity during cognitive tasks [78]? Given that the two modalities become less correlated in transmodal cortex in the resting state, the relationship between them during task may depend on demand and cognitive functions required to complete the task.

Finally, the present results should be interpreted in light of several methodological considerations. First, although we conduct extensive sensitivity testing, including multiple ways of defining functional connectivity, there exist many more ways in the literature to estimate both fMRI and MEG connectivity [100, 143]. Second, to ensure that the analyses were performed in the same participants using both resting state fMRI and MEG, and that the participants have no familial relationships, we utilized a reduced version of the HCP sample. Third, in order to directly compare the contributions of multiple frequency bands, all were entered into the same model. As a result however, the observations in the linear models (network edges) are not independent, violating a basic assumption of these statistical models. For this reason, we only use model fits and dominance values to compare the correspondence of fMRI and MEG across a set of nodes, each of which is estimated under the same conditions. Finally, to ensure that the findings are independent from sensitivity of MEG to neural activity from different regions, we compared the cross-modal correspondence map with MEG signal-to-noise ratio and source localization error, where no significant associations were identified. However, MEG is still susceptible to such artifacts given that regions with lower signal-to-noise ratio (e.g. Sylvian fissure) are the ones where source reconstruction solutions have higher source localization errors [53, 66].

Despite complementary strengths to image spatiotemporal brain dynamics, the links between MEG and fMRI are not fully understood and the two fields have diverged. The present report bridges the two disciplines by comprehensively mapping haemodynamic and electromagnetic network architectures. By considering the contributions of the canonical frequency bands simultaneously, we show that the superposition and mixing of MEG neurophysiological rhythms manifests as highly structured patterns of fMRI functional connectivity. Systematic convergence and divergence among the two modalities in different brain regions opens fundamentally new questions about the relationship between cortical hierarchies and multi-modal functional networks.

## METHODS

### Dataset: Human Connectome Project (HCP)

Resting state magnetoencephalography (MEG) data of a set of healthy young adults (*n* = 33; age range 22-35 years) with no familial relationships were obtained from Human Connectome Project (HCP; S900 release [140]). The data includes resting state scans of about 6 minutes long (sampling rate = 2034.5 Hz; anti-aliasing filter lowpass filter at 400 Hz) and noise recordings for all participants. MEG anatomical data and 3T structural magnetic resonance imaging (MRI) data of all participants were also obtained for MEG pre-processing. Finally, we obtained functional MRI data of the same *n* = 33 individuals from HCP dataset. All four resting state fMRI scans (two scans with R/L and L/R phase encoding directions on day 1 and day 2, each about 15 minutes long; *TR* = 720 ms) were available for all participants.

### HCP Data Processing

#### Resting state magnetoencephalography (MEG)

Resting state MEG data was analyzed using Brainstorm software, which is documented and freely available for download online under the GNU general public license ([133]; http://neuroimage.usc.edu/brainstorm). The MEG recordings were registered to the structural MRI scan of each individual using the anatomical transformation matrix provided by HCP for co-registration, following the procedure described in Brainstorm’s online tutorials for the HCP dataset (https://neuroimage.usc.edu/brainstorm/Tutorials/HCP-MEG). The pre-processing was performed by applying notch filters at 60, 120, 180, 240, and 300 Hz, and was followed by a highpass filter at 0.3 Hz to remove slow-wave and DC-offset artifacts. Bad channels were marked based on the information obtained through the data management platform of HCP for MEG data (ConnectomeDB; https://db.humanconnectome.org/). The artifacts (including heartbeats, eye blinks, saccades, muscle movements, and noisy segments) were then removed from the recordings using automatic procedures as proposed by Brainstorm. More specifically, electrocardiogram (ECG) and electrooculogram (EOG) recordings were used to detect heartbeats and blinks, respectively. We then used Signal-Space Projections (SSP) to automatically remove the detected artifacts. We also used SSP to remove saccades and muscle activity as low-frequency (1-7 Hz) and high-frequency (40-240 Hz) components, respectively.

The pre-processed sensor-level data was then used to obtain a source estimation on HCP’s fsLR4k cortex surface for each participant. Head models were computed using overlapping spheres and the data and noise covariance matrices were estimated from the resting state MEG and noise recordings. Linearly constrained minimum variance (LCMV) beamformers method from Brainstorm was then used to obtain the source activity for each participant. We performed data covariance regularization and normalized the estimated source variance by the noise covariance matrix to reduce the effect of variable source depth. The L2 matrix norm (i.e. regularization parameter) of data covariance matrix is usually defined as the largest eigenvalue of its eigenspectrum. However, the eigenspectrum of MEG data covariance can be ill-conditioned, such that the eigenvalues may span many decades where larger eigenvalues are overestimated and smaller eigenvalues are underestimated. In other words, the L2 norm of the data covariance matrix can be many times larger than the majority of eigenvalues, making it difficult to select a conventional regularization parameter. Following guidelines from Brainstorm [133], we used the “median eigenvalue” method to regularize the data covariance matrix, where the eigenvalues smaller than the median eigenvalue are replaced with the median eigenvalue itself (i.e. flattening the tail of eigenvalues spectrum to the median). The covariance matrix is then reconstructed using the modified eigenspectrum. This helps to avoid the instability of data covariance inversion caused by the smallest eigenvalues and regularizes the data covariance matrix. Source orientations were constrained to be normal to the cortical surface at each of the 8,000 vertex locations on the fsLR4k surface. Source-level time-series were then parcellated into 400 regions using the Schaefer-400 atlas [119], such that a given parcel’s time series was estimated as the first principal component of its constituting sources’ time series.

Parcellated time-series were then used to estimate functional connectivity with an amplitude-based connectivity measure from Brainstorm (amplitude envelope correlation; AEC [20]). An orthogonalization process was applied to correct for the spatial leakage effect by removing all shared zero-lag signals [28]. AEC functional connectivity were derived for each participant at six canonical electrophysiological bands (i.e., delta (*δ*: 2-4 Hz), theta (*θ*: 5-7 Hz), alpha (*α*: 8-12 Hz), beta (*β*: 15-29 Hz), low gamma (lo-*γ*: 30-59 Hz), and high gamma (hi-*γ*: 60-90Hz)). Group-average MEG functional connectivity matrices were constructed as the mean functional connectivity across all individuals for each frequency band. For comparison, band-limited group-average AEC matrices were also estimated without correcting for spatial leakage effect.

We also processed the MEG data using additional methodological choices. First, the LCMV source reconstructed and parcellated time-series were used to estimate functional connectivity with an alternative, phase-based connectivity measure (phase locking value; PLV [79, 96]) for each frequency band. Second, another source reconstruction method (standardized low resolution brain electromagnetic tomography; sLoreta [105]) was used instead of LCMV beamformers to obtain source-level time-series, given that previous reports suggest that sLoreta improves source localization accuracy [65, 67]. Source-level time-series, obtained by sLoreta, were then parcellated into 400 regions and were used to estimate AEC matrices with spatial leakage correction for the six frequency bands. Third, to ensure that the findings are independent from choice of parcellation, a low resolution atlas (Schaefer-200 [119]) was used to parcellate the original LCMV source-level time-series to 200 cortical regions and obtain spatial leakage corrected AEC connectivity matrices. Finally, we estimated MEG source localization errors for LCMV and sLoreta source reconstruction solutions using cross-talk functions (CTF) [64– 67, 82, 95]. CTF of a given source *i* is a measure of how activity from all other sources contributes to the activity estimated for the *i*-th source. Following guidelines from Brainstorm [133] and MNE-Python software packages [56], we used CTF to calculate peak localization error of a given source *i* as the Euclidean distance between the peak location estimated for source *i* and the true source location *i* on the surface model [65, 95]. Source-level CTF was then parcellated using the Schaefer-400 atlas. We also estimated source-level signal-to-noise ratio (SNR) for LCMV source reconstruction solution as follows [53, 106]:

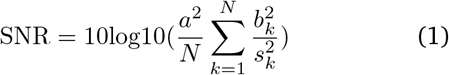

where *a* is the source amplitude (i.e. typical strength of a dipole, which is 10 nAm [58]), *N* is the number of sensors, *b*_*k*_ is the signal at sensor *k* estimated by the forward model for a source with unit amplitude, and 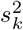 is the noise variance at sensor *k*. SNR was first calculated at the source level and was then parcellated using the Schaefer-400 atlas.

#### Resting state functional MRI

The functional MRI data were pre-processed using HCP minimal pre-processing pipelines [52, 140]. Detailed information regarding data acquisition and preprocessing is available elsewhere [52, 140]. Briefly, all 3T functional MRI time-series (voxel resolution of 2 mm isotropic) were corrected for gradient nonlinearity, head motion using a rigid body transformation, and geometric distortions using scan pairs with opposite phase encoding directions (R/L, L/R) [34]. Further pre-processing steps include co-registration of the corrected images to the T1w structural MR images, brain extraction, normalization of whole brain intensity, high-pass filtering (*>* 2000s FWHM; to correct for scanner drifts), and removing additional noise using the ICA-FIX process [34, 116]. The pre-processed time-series were then parcellated into 400 cortical areas using Schaefer-400 parcellation [119]. The parcellated time-series were used to construct functional connectivity matrices as Pearson correlation coefficients between pairs of regional time-series for each of the four scans and each participant. A group-average functional connectivity matrix was constructed as the mean functional connectivity across all individuals and scans.

#### Diffusion weighted imaging (DWI)

Diffusion weighted imaging (DWI) data was obtained for the same individuals from the HCP dataset. MRtrix3 package [137] (https://www.mrtrix.org/) was used to pre-process the DWI data as described elsewhere [124]. In brief, multi-shell multi-tissue constrained spherical deconvolution algorithm from MRtrix was applied to generate fiber orientation distributions [37, 77]. Probabilistic streamline tractography based on the generated fiber orientation distributions was used to reconstruct white matter edges [136]. The tract weights were optimized by estimating an appropriate cross-section multiplier for each streamline following the procedure proposed by Smith and colleagues [128]. Structural connectivity matrices were then reconstructed for each participant using the Schaefer-400 atlas [119]. Finally, a binary group-level structural connectivity matrix was constructed using a consensus approach that preserves the edge length distribution in individual participants [14, 94]. The binary consensus structural connectivity matrix was weighted by the average structural connectivity across individuals to obtain a weighted structural connectivity matrix.

### BigBrain histological data

To characterize the cytoarchitectural variation across the cortex, cell-staining intensity profile data were obtained from the BigBrain atlas [2, 104]. The BigBrain is a high-resolution (20 *μm*) histological atlas of a post mortem human brain and includes cell-staining intensities that are sampled at each vertex across 50 equivolumetric surfaces from the pial to the white matter surface using the Merker staining technique [2, 92]. The staining intensity profile data represent neuronal density and soma size at varying cortical depths, capturing the regional differentiation of cytoarchitecture [2, 103, 104, 144, 145]. Intensity profiles at various cortical depths can be used to approximately identify boundaries of cortical layers that separate supragranular (cortical layers I-III), granular (cortical layer IV), and infragranular (cortical layers V-VI) layers [104, 144, 145]. The data were obtained on *fsaverage* surface (164k vertices) from the BigBrainWarp toolbox [104] and were parcellated into 400 cortical regions using the Schaefer-400 atlas [119].

The cross-modal correspondence map, estimated as adjusted-*R*^2^ (See “Multi-linear model” for more details), was then compared with the parcellated cell-staining intensity data. Specifically, the regional model fit was correlated with cell-staining profiles at each cortical depth using Spearman rank correlation (*r*_*s*_). 10,000 spatialautocorrelation preserving nulls were used to construct a null distribution of correlation at each cortical depth (See “Null model” for more details on spatial-autocorrelation preserving nulls). Significance of the associations were estimated by comparing the empirical Spearman rank correlation with the distribution of null correlations at each cortical depth, identifying the number of null correlations that were equal to or greater than the empirical correlation (two-tailed test). Finally, Benjamini-Hochberg procedure [13] was used to correct for multiple comparisons by controlling the false discovery rate (FDR) at 5% across all 50 comparisons.

### Allen Human Brain Atlas (AHBA)

Regional microarray expression data were obtained from 6 post-mortem brains (1 female, ages 24.0–57.0,42.50 ± 13.38) provided by the Allen Human Brain Atlas (AHBA, https://human.brain-map.org; [69]). Data were processed with the abagen toolbox (version 0.1.3-doc; https://github.com/rmarkello/abagen; [88]) using the Schaefer-400 volumetric atlas in MNI space [119].

First, microarray probes were reannotated using data provided by [3]; probes not matched to a valid Entrez ID were discarded. Next, probes were filtered based on their expression intensity relative to background noise [109], such that probes with intensity less than the background in ≥50.00% of samples across donors were discarded. When multiple probes indexed the expression of the same gene, we selected and used the probe with the most consistent pattern of regional variation across donors (i.e., differential stability; [68]), calculated with:

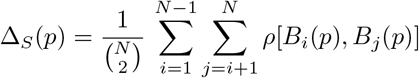

where *ρ* is Spearman’s rank correlation of the expression of a single probe, p, across regions in two donors *B*_*i*_ and *B*_*j*_, and N is the total number of donors. Here, regions correspond to the structural designations provided in the ontology from the AHBA.

The MNI coordinates of tissue samples were updated to those generated via non-linear registration using the Advanced Normalization Tools (ANTs; https://github.com/chrisfilo/alleninf). To increase spatial coverage, tissue samples were mirrored bilaterally across the left and right hemispheres [111]. Samples were assigned to brain regions in the provided atlas if their MNI coordinates were within 2 mm of a given parcel. If a brain region was not assigned a tissue sample based on the above procedure, every voxel in the region was mapped to the nearest tissue sample from the donor in order to generate a dense, interpolated expression map. The average of these expression values was taken across all voxels in the region, weighted by the distance between each voxel and the sample mapped to it, in order to obtain an estimate of the parcellated expression values for the missing region. All tissue samples not assigned to a brain region in the provided atlas were discarded.

Inter-subject variation was addressed by normalizing tissue sample expression values across genes using a robust sigmoid function [48]:

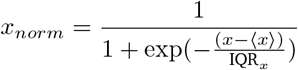

where ⟨*x*⟩ is the median and IQR_*x*_ is the normalized interquartile range of the expression of a single tissue sample across genes. Normalized expression values were then rescaled to the unit interval:

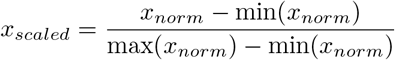

Gene expression values were then normalized across tissue samples using an identical procedure. Samples assigned to the same brain region were averaged separately for each donor and then across donors, yielding a regional expression matrix of 15,633 genes. Expression of NPY1R (Neuropeptide Y Receptor Y1) was extracted from the regional expression matrix and was related to the cross-modal correspondence map, estimated as adjusted-*R*^2^ (See “Multi-linear model” for more details), using 10,000 spatial-autocorrelation preserving nulls (See “Null models” for more details).

### Multi-linear model

#### Regional model

A multiple linear regression model was used to assess regional associations between haemodynamic (fMRI) and electromagnetic (MEG) functional connectivity (Fig. 1 [142]). A separate multi-linear model is applied for each brain region from the parcellated data, predicting the region’s fMRI functional connectivity profile from its band-limited MEG functional connectivity. The dependent variable is a row of the fMRI connectivity matrix and the independent variables (predictors) are the corresponding rows of MEG connectivity for the six canonical electrophysiological bands. The linear regression model for each brain region *i* is constructed as follows:

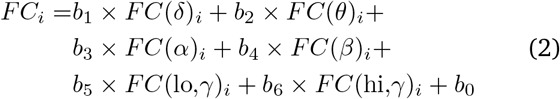

where the dependant variable *FC*_*i*_ is the set of fMRI connections of node *i* to the other *j* ≠ *i* regions and the predictors are sets of MEG connections of node *i* to the other *j* ≠ *i* regions for the six frequency bands (*FC*(*δ*)_*i*_, *FC*(*θ*)_*i*_, *FC*(*α*)_*i*_, *FC*(*β*)_*i*_, *FC*(lo,*γ*)_*i*_, *FC*(hi,*γ*)_*i*_). The regression coefficients *b*_1_, …, *b*_6_ and the intercept *b*_0_ are then optimized to yield maximum correlation between empirical and predicted fMRI connectivity for each brain region. Goodness of fit for each regional model is quantified using adjusted-*R*^2^ (coefficient of determination).

#### Global model

For comparison with the regional model, a single global model was fitted to the data, predicting the wholebrain fMRI functional connectivity from the whole-brain band-limited MEG functional connectivity (Fig. 1d). Specifically, rather than applying a multi-linear model for each region (i.e. each row) separately, we fit a single multi-linear model using the upper triangle of bandlimited MEG connectivity (i.e. all values above the diagonal of MEG connectivity matrices) as predictors and predict the upper triangle of fMRI connectivity. The equation below describes the multi-linear global model:

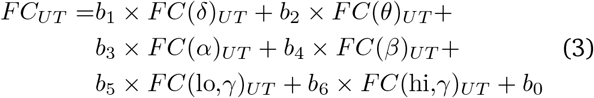

where the dependent variable *FC*_*UT*_ is the vectorized upper triangle of fMRI functional connectivity (i.e. above diagonal values) and the predictors are the vectorized upper triangles of MEG functional connectivity for the six frequency bands. The regression coefficients *b*_1_, …, *b*_6_ and the intercept *b*_0_ are then optimized to yield maximum correlation between empirical and predicted fMRI connectivity. Similar to the regional model, the goodness of fit for the global model is quantified using adjusted-*R*^2^ (coefficient of determination).

#### Region-level cross-validation

Region-level cross-validation was performed to assess out-of-sample model performance. Given the spatial autocorrelation inherent to the data, random splits of brain regions into train and test sets may result in out-of-sample correlations that are inflated due to spatial proximity [90]. To take this into account, we used a distance-dependant cross-validation approach where we pseudorandomly split the connectivity profile of a given region (e.g. node *i*) into train and test sets based on spatial separation [59]. We used inter-regional Euclidean distance to select 75% of the closest regions to a randomly selected source region as the train set and the remaining 25% of the regions as test set. The random source region can be any of the 399 regions connected to node *i*; hence, the connectivity profile of node *i* is split into 399 unique train and test sets. We then train the multi-linear model using the train set and predict functional connectivity of the test set for each region and each split. Finally, the model performance is quantified using Pearson correlation coefficient between empirical and predicted values. The cross-validated regional model performance is then estimated as the mean correlation coefficient between empirical and predicted values across splits for each brain region.

#### Subject-level cross-validation

Leave-one-out cross-validation was performed to assess model performance on held-out subjects. Briefly, the regional multi-linear model is trained using the group-average data from *n* −1 subjects. The trained model is then used to predict fMRI connectivity profile of each region on the held-out subject (test set). The model performance is quantified as the Pearson correlation coefficient between empirical and predicted connectivity of each region. The analysis is repeated for all subjects and the regional model performance is averaged across individuals.

### Diffusion map embedding

Diffusion map embedding was used to identify the principal axis of variation in functional organization of the cortex (diffusion map embedding and alignment package; https://github.com/satra/mapalign) [80, 87]. Diffusion map embedding is a nonlinear dimensionality reduction technique that generates a low-dimensional representation of high-dimensional data by projecting it into an embedding space, such that the areas with similar connectivity profiles will be closer in distance in the new common space compared to the areas with dissimilar connectivity profiles [27, 80, 87]. In brief, following the procedure described by Margulies and colleagues [87], each row of the group-average fMRI functional connectivity was thresholded at 90%, such that only the top 10% of functional connections was retained in the matrix. Next, a cosine-similarity matrix was estimated based on the remaining functional connections, where the resulting pairwise cosine distances represent the similarity between the connectivity profiles of cortical regions according to their strongest connections. Finally, the diffusion map embedding was applied to the resulting positive affinity matrix. This identifies the principal axis of variation in functional connectivity, along which cortical regions are ordered based on the similarity of their connectivity profiles. The identified functional gradient or hierarchy spans the unimodal-transmodal axis, separating primary sensory-motor cortices from association cortex. The functional gradient map is also available as part of the neuromaps toolbox [89]. The functional gradient was used as a metric of hierarchical organization of the cortex and was compared with the regional model fit (Fig. 2).

### Structure-function coupling

Structure-function coupling was estimated following the procedure described by Baum and colleagues [11]. Structural and functional connectivity profiles of each brain region (i.e. each row of the connectivity matrices) were extracted from the weighted group-level structural and functional connectivity matrices. Structurefunction coupling of a given region was then estimated as the Spearman rank correlation between non-zero values of that region’s structural and functional connectivity profiles. Finally, the resulting whole-brain structure-function coupling map was compared with the cross-modal correspondence map (i.e. *R*^2^ map from the regional model). Significance of the association between the two maps was assessed using 10,000 spatial-autocorrelation preserving nulls (See “Null model” for more details).

### Dominance analysis

Dominance Analysis was used to quantify the distinct contributions of resting state MEG connectivity at different frequency bands to the prediction of resting state fMRI connectivity in the multi-linear model [4, 21] (https://github.com/dominance-analysis/dominance-analysis). Dominance analysis estimates the relative importance of predictors by constructing all possible combinations of predictors and re-fitting the multilinear model for each combination (a model with *p* predictors will have 2^*p*^ −1 models for all possible combinations of predictors). The relative contribution of each predictor is then quantified as increase in variance explained by adding that predictor to the models (i.e. gain in adjusted-*R*^2^). Here we first constructed a multiple linear regression model for each region with MEG connectivity profile of that region at six frequency bands as independent variables (predictors) and fMRI connectivity of the region as the dependent variable to quantify the distinct contribution of each factor using dominance analysis. The relative importance of each factor is estimated as “percent relative importance”, which is a summary measure that quantifies the percent value of the additional contribution of that predictor to all subset models.

### Null model

To make inferences about the topographic correlations between any two brain maps, we implement a null model that systematically disrupts the relationship between two topographic maps but preserves their spatial autocorrelation [1, 90]. We used the Schaefer-400 atlas in the HCP’s fsLR32k grayordinate space [119, 140]. The spherical projection of the fsLR32k surface was used to define spatial coordinates for each parcel by selecting the vertex closest to the center-of-mass of each parcel [125, 141, 142]. The resulting spatial coordinates were used to generate null models by applying randomly-sampled rotations and reassigning node values based on the closest resulting parcel (10,000 repetitions). The rotation was applied to one hemisphere and then mirrored to the other hemisphere.

### Code and data availability

Code used to conduct the reported analyses is available on GitHub (https://github.com/netneurolab/shafiei_megfmrimapping). Data used in this study were obtained from the Human Connectome Project (HCP) database (available at https://db.humanconnectome.org/). The data and code needed to generate all main and supplementary figures can be found in https://github.com/netneurolab/shafiei_megfmrimapping and https://zenodo.org/record/6728338.

## ACKNOWLEDGMENTS

We thank Ross Markello, Estefany Suarez, Bertha Vazquez-Rodriguez, Zhen-Qi Liu for their comments on the manuscript. This research was undertaken thanks in part to funding from the Canada First Research Excellence Fund, awarded to McGill University for the Healthy Brains for Healthy Lives initiative. BM acknowledges support from the Natural Sciences and Engineering Research Council of Canada (NSERC Discovery Grant RGPIN #017-04265) and from the Canada Research Chairs Program. SB is grateful for the support received from the NIH (R01 EB026299), a Discovery grant from the Natural Science and Engineering Research Council of Canada (NSERC 436355-13), the CIHR Canada research Chair in Neural Dynamics of Brain Systems, the Brain Canada Foundation with support from Health Canada, and the Innovative Ideas program from the Canada First Research Excellence Fund, awarded to McGill University for the Healthy Brains for Healthy Lives initiative. GS acknowledges support from the Natural Sciences and Engineering Research Council of Canada (NSERC) and the Fonds de recherche du Québec - Nature et Technologies (FRQNT).

---

**Table S1.** **Analysis of Variance (ANOVA) for dominance analysis** | To quantitatively assess the differences in band-specific contributions to the cross-modal correspondence map, contributions estimated from dominance analysis were compared for all possible pairs of frequency bands using Analysis of Variance (ANOVA). All reported *p*-values are from two-tailed tests and are corrected for multiple comparisons using Bonferroni correction. Cohen’s *d* denotes effect size.

| Band A | Band B | mean(A) | mean(B) | difference | *t*-value | *p*-value | Cohen's *d* |
| --- | --- | --- | --- | --- | --- | --- | --- |
| delta | theta | 15.06 | 19.87 | -4.81 | -5.26 | <0.0001 | -0.37 |
| delta | alpha | 15.06 | 18.79 | -3.72 | -4.07 | 0.00073 | -0.29 |
| delta | beta | 15.06 | 27.44 | -12.37 | -13.53 | <0.0001 | -0.96 |
| delta | lo-gamma | 15.06 | 11.28 | 3.78 | 4.14 | 0.00055 | 0.29 |
| delta | hi-gamma | 15.06 | 7.56 | 7.51 | 8.21 | <0.0001 | 0.58 |
| theta | alpha | 19.87 | 18.79 | 1.08 | 1.19 | 1 | 0.08 |
| theta | beta | 19.87 | 27.44 | -7.56 | -8.27 | <0.0001 | -0.58 |
| theta | lo-gamma | 19.87 | 11.28 | 8.59 | 9.39 | <0.0001 | 0.66 |
| theta | hi-gamma | 19.87 | 7.56 | 12.32 | 13.46 | <0.0001 | 0.95 |
| alpha | beta | 18.79 | 27.44 | -8.65 | -9.46 | <0.0001 | -0.69 |
| alpha | lo-gamma | 18.79 | 11.28 | 7.51 | 8.21 | <0.0001 | 0.58 |
| alpha | hi-gamma | 18.79 | 7.56 | 11.23 | 12.28 | <0.0001 | 0.87 |
| beta | lo-gamma | 27.44 | 11.28 | 16.16 | 17.66 | <0.0001 | 1.25 |
| beta | hi-gamma | 27.44 | 7.56 | 19.88 | 21.73 | <0.0001 | 1.54 |
| lo-gamma | hi-gamma | 11.28 | 7.56 | 3.72 | 4.07 | 0.00072 | 0.29 |

**Figure S1.**
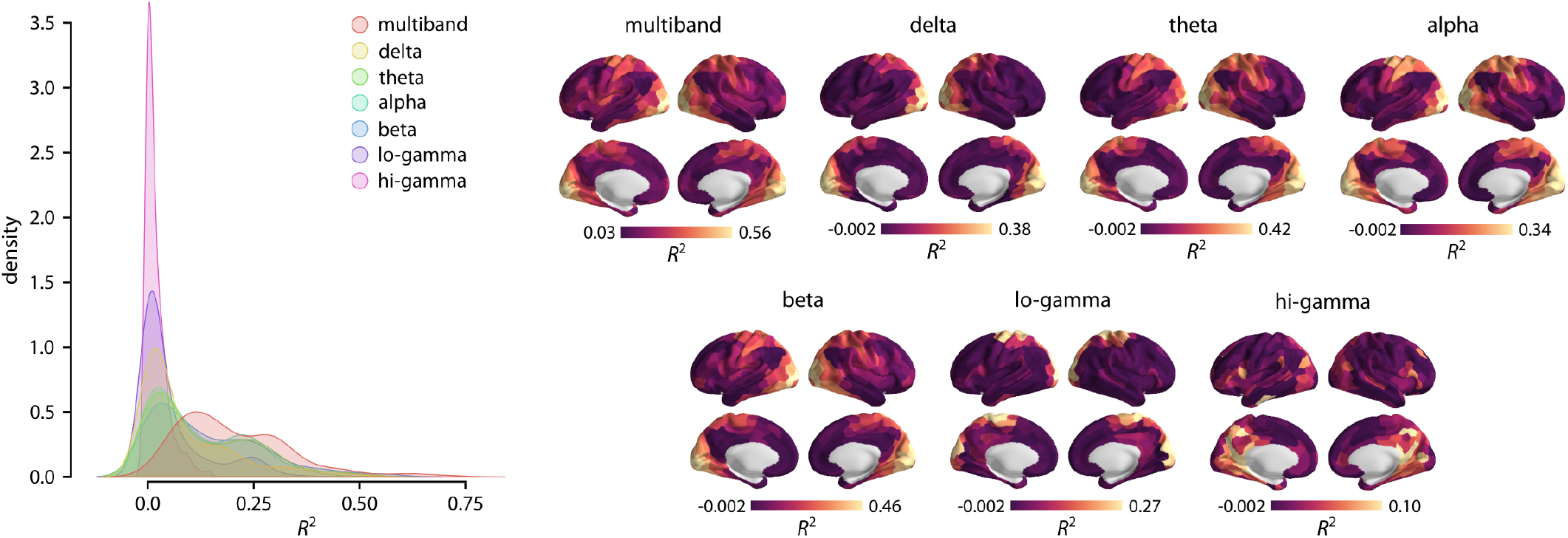
Band-specific regional model fit. Separate regional regression models were applied to map MEG functional connectivity (AEC) to fMRI functional connectivity at each frequency band. Distributions of adjusted-*R*^2^ are depicted for band-specific regional model fits and for the multiband model fit obtained by the original analysis. The multi-linear regional model that combines MEG connectivity at multiple rhythms to predict regional fMRI connectivity profiles performs better than the band-specific models. The data and code needed to generate this figure can be found in https://github.com/netneurolab/shafiei_megfmrimapping and https://zenodo.org/record/6728338.

**Figure S2.**
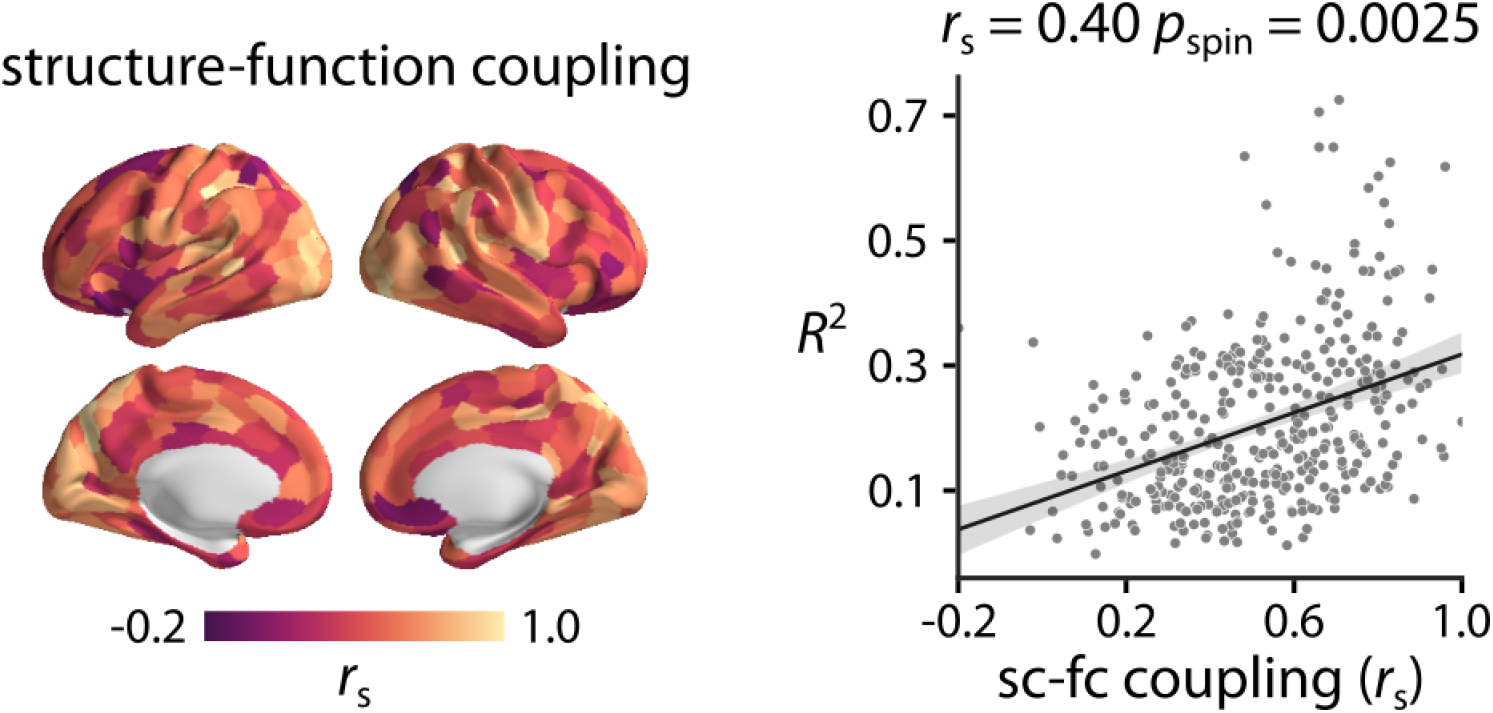
Structure-function coupling. Structure-function coupling was estimated as the Spearman rank correlation (*r*_*s*_) between regional structural and functional connectivity profiles [11]. The cross-modal *R*^2^ map (i.e. regional model fit) is then compared with the structure-function coupling across the cortex. The data and code needed to generate this figure can be found in https://github.com/netneurolab/shafiei_megfmrimapping and https://zenodo.org/record/6728338.

**Figure S3.**
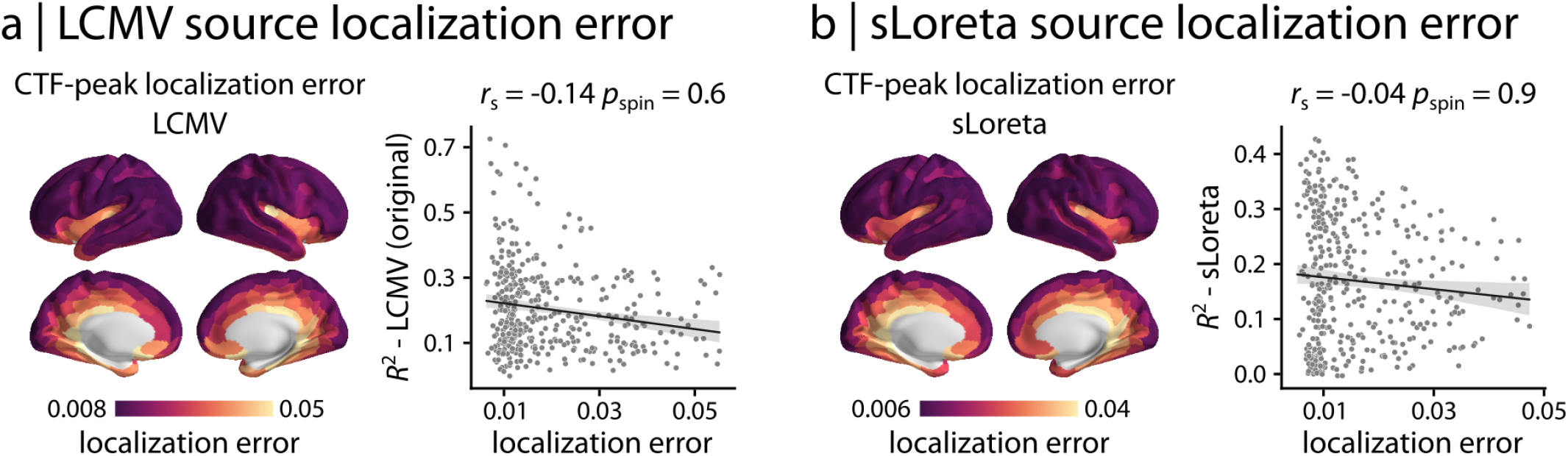
Source localization error. MEG source localization error is estimated for (a) LCMV and (b) sLoreta source reconstruction solutions using cross-talk functions (CTF) [64, 65, 67, 82, 95]. CTF is used to calculate peak localization error of a given source *i* as the Euclidean distance between the peak location estimated for source *i* and the true source location *i* on the surface model [65, 95]. No significant association is observed between the cross-modal correspondence *R*^2^ map and peak localization error for LCMV and sLoreta. The data and code needed to generate this figure can be found in https://github.com/netneurolab/shafiei_megfmrimapping and https://zenodo.org/record/6728338.

**Figure S4.**
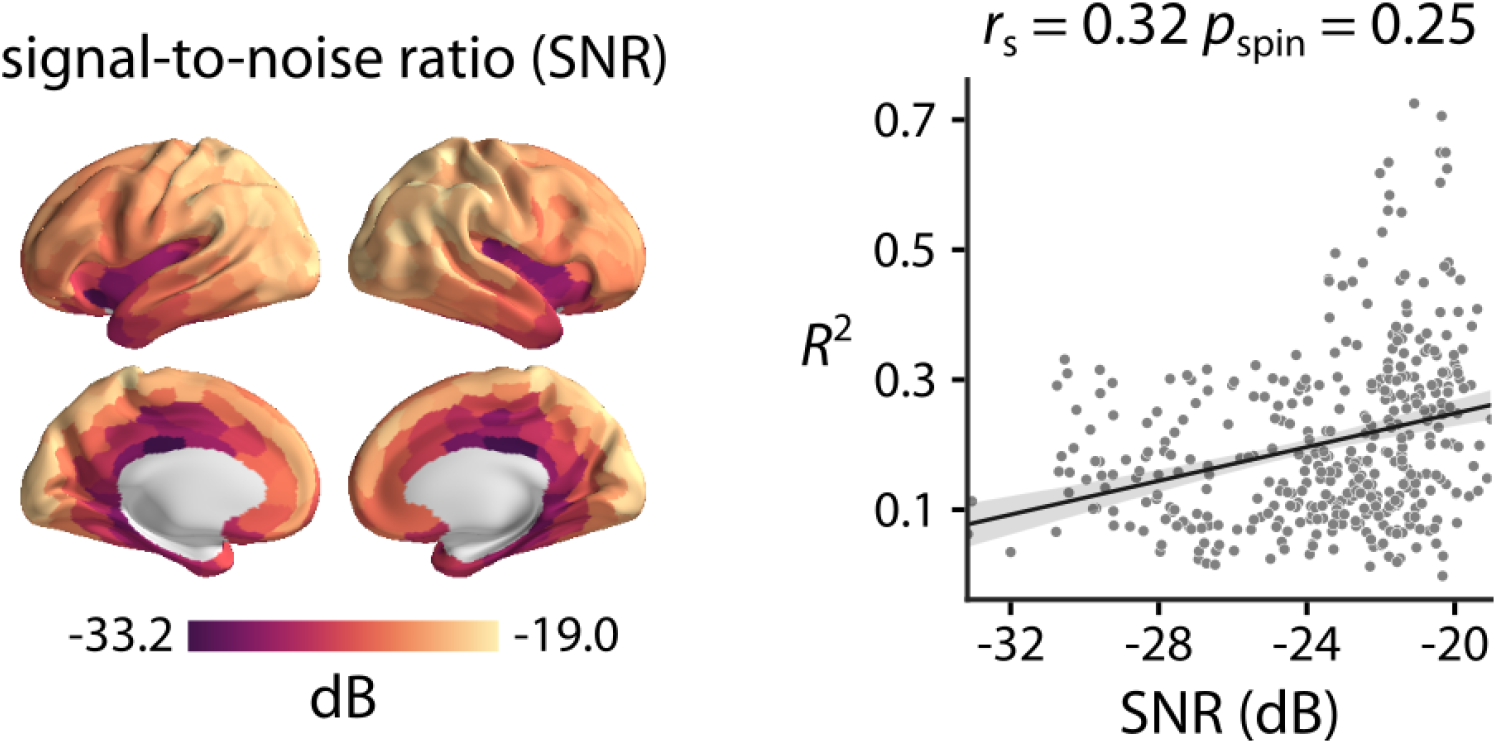
Signal-to-noise ratio. MEG signal-to-noise ratio (SNR) was estimated at the source level. Parcellated, group-average SNR map is depicted across the cortex. The cross-modal correspondence *R*^2^ map (i.e. regional model fit) is then compared with the SNR map. The data and code needed to generate this figure can be found in https://github.com/netneurolab/shafiei_megfmrimapping and https://zenodo.org/record/6728338.

**Figure S5.**
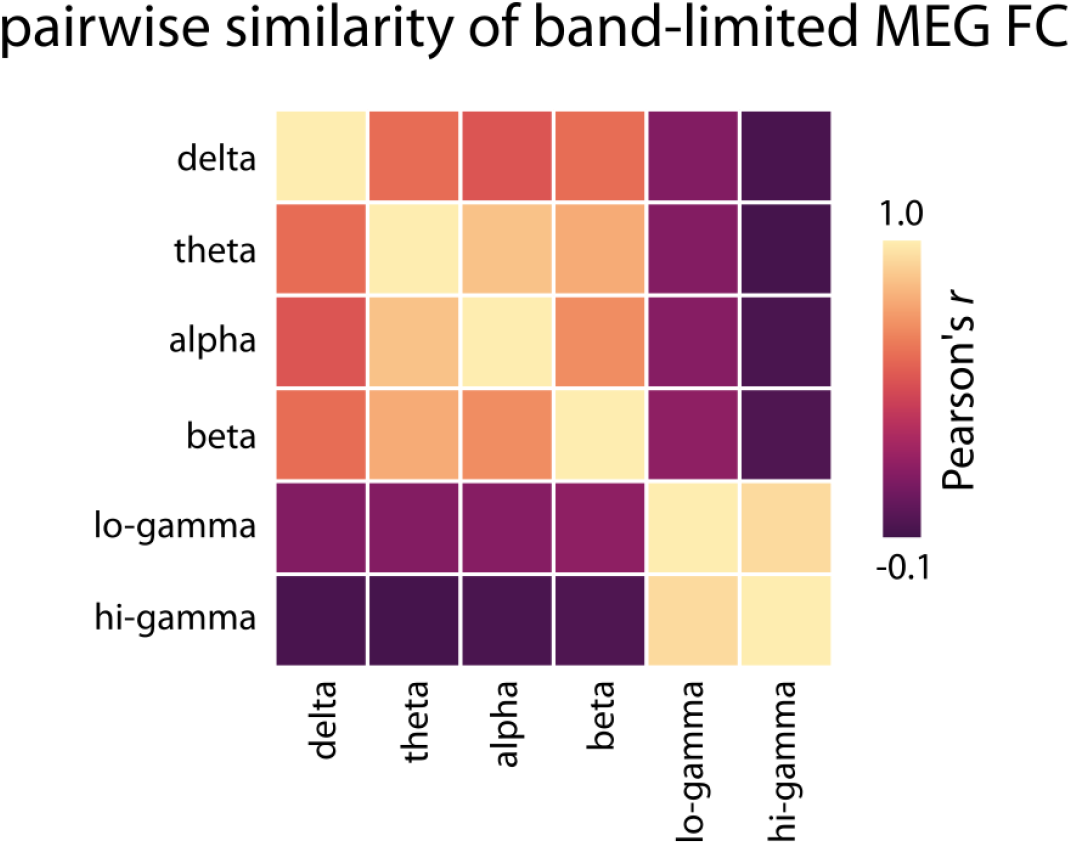
Pairwise similarity of band-limited MEG functional connectivity. Pearson correlation coefficient is calculated between upper triangles (i.e. values above diagonal) of band-limited MEG AEC functional connectivity to assess the pairwise similarity between MEG connectivity maps. The data and code needed to generate this figure can be found in https://github.com/netneurolab/shafiei_megfmrimapping and https://zenodo.org/record/6728338.

